# Intravital imaging of cerebral microinfarct reveals an astrocyte reaction led to glial scar

**DOI:** 10.1101/2021.09.29.462492

**Authors:** Jingu Lee, Joon-Goon Kim, Sujung Hong, Young Seo Kim, Soyeon Ahn, Ryul Kim, Heejung Chun, Ki Duk Park, Yong Jeong, Dong-Eog Kim, C. Justin Lee, Taeyun Ku, Pilhan Kim

**Affiliations:** Graduate School of Nanoscience and Technology, Korea Advanced Institute of Science and Technology (KAIST), Daejeon, Republic of Korea; KI for Health Science and Technology (KIHST), Korea Advanced Institute of Science and Technology (KAIST), Daejeon, Republic of Korea; Graduate School of Medical Science and Engineering, Korea Advanced Institute of Science and Technology (KAIST), Daejeon, Republic of Korea; Division of Hematology-Oncology, Department of Medicine, Samsung Medical Center, Sungkyunkwan University School of Medicine, Seoul, Republic of Korea; Center for Cognition and Sociality, Institute for Basic Science (IBS), Daejeon, Republic of Korea; Convergence Research Center for Diagnosis, Treatment and Care System of Dementia, Korea Institute of Science and Technology (KIST), Seoul, Republic of Korea; Division of Bio-Med Science & Technology, KIST School, University of Science and Technology (UST), Seoul; Department of Bio and Brain Engineering, Korea Advanced Institute of Science and Technology (KAIST), Daejeon, Republic of Korea; Department of Neurology, Dongguk University College of Medicine, Dongguk University Ilsan Hospital, Goyang, Korea

**Keywords:** Cerebral microinfarct, Glial scar formation, Intravital imaging, Reactive astrocyte, Blood-brain barrier

## Abstract

Cerebral microinfarct increases the risk of dementia. But how microscopic cerebrovascular disruption affects the brain tissue in cellular-level are mostly unknown. Herein, with a longitudinal intravital imaging, we serially visualized in vivo dynamic cellular-level changes in astrocyte, pericyte and neuron as well as microvascular integrity after the induction of cerebral microinfarction for 1 month in mice. At day 2-3, it revealed a localized edema with acute astrocyte loss, neuronal death, impaired pericyte-vessel coverage and extravascular leakage indicating blood-brain barrier (BBB) dysfunction. At day 5, edema disappeared with recovery of pericyte-vessel coverage and BBB integrity. But brain tissue continued to shrink with persisted loss of astrocyte and neuron in microinfarct until 30 days, resulting in a collagen-rich fibrous scar surrounding the microinfarct. Notably, reactive astrocytes appeared at the peri-infarct area early at day 2 and thereafter accumulated in the peri-infarct. Oral administration of a reversible monoamine oxidase B inhibitor significantly decreased the astrocyte reactivity and fibrous scar formation. Our result suggests that astrocyte reactivity may be a key target to alleviate the impact of microinfarction.

## Introduction

Cerebral microinfarcts, small ischemic lesions, can be caused by cerebral small vessel disease, microemboli, and hypoperfusion(van Veluw, Shih et al., 2017). Prevalent microinfarcts contribute to the disruption of structural brain connection, which can progress to a cognitive impairment and dementia that is independent of Alzheimer’s disease (AD) pathology(Smith, Schneider et al., 2012, van Veluw et al., 2017). A systemic review reported that the cerebral microinfarct prevalence was 62% in vascular dementia patient, 43% in AD patient, and 24% in healthy aged individuals(Brundel, de Bresser et al., 2012). With a growing body of evidence of an association between cerebral microinfarcts and dementia, there are increasing studies to improve the understanding of the microinfarct as a risk factor and its functional impact(Freeze, Bacskai et al., 2019, Okamoto, Yamamoto et al., 2012, Shih, Hyacinth et al., 2018). It has been shown that cerebrovascular dysfunction could cause spontaneous thrombotic cerebral microinfarcts, followed by cerebral amyloid angiopathy (CAA), blood-brain barrier (BBB) breakdown, and cognitive impairment(Shih, Blinder et al., 2013, Tan, Xue et al., 2015). However, little is known about the mechanistic link between microscopic cerebrovascular disruption and brain damage. Furthermore, dynamic *in vivo* cellular-level changes in glial, vascular and neuronal cells after the onset of cerebral microinfarction have been mostly unknown partly due to technical difficulties in the longitudinal high-resolution *in situ* observation of their complex behaviors and fates serially in a live animal models *in vivo* over long period of time.

In this study, we have achieved a longitudinal cellular level intravital imaging of a cerebral microinfarction mouse model to visualize highly dynamic behaviors of astrocytes and pericytes and changes in vascular obstruction and leakage for 1 month. The intravital imaging of the microinfarction lesion revealed an acute tissue expansion, localized edema, at day 2 followed by gradual tissue shrinkage for 1 month, resulting in a cerebral atrophy with a neuronal loss and collagen-rich scar formation surrounding the microinfarct. An acute astrocyte loss and neuronal death observed after the microinfarct induction was persisted until 1 month whereas acute impairment of pericyte-vessel coverage, microvascular flow and vascular leakage of 3 kDa dextran (but not 2 MDa dextran) were partially recovered at day 5. Meanwhile, glial fibrillary acidic protein (GFAP)-expressing reactive astrocytes were highly accumulated at the peri-infarct area for 1 month after the microinfarct induction. Thus, we further investigated the role of GFAP^+^ reactive astrocytes as a driver of fibrous scar formation associated with the cerebral microinfarct. GFAP^+^ reactive astrocytes started to be observed at the peri-infarct area from 2 days after the microinfarct induction and appeared to be accumulated with glial scar formation until 1 month. Additionally, GFAP^+^ astrocytes co-expressing lipocalin2 (LCN2), severe reactive astrocytes, were observed at the peri-infarct area with glial scar formation in the infarct. Oral administration of a reversible monoamine oxidase B (MAO-B) inhibitor, KDS2010, to alleviate astrocytes reactivity significantly decreased the collagen-rich glial scar formation after the microinfarct induction.

## Results

### Glial fibrous scar formation in cerebral microinfarct

A microinfarction in a penetrating arteriole identified by *in vivo* CD31 labeling of endothelial cells (EC) in cerebral cortex was induced by photothrombosis (PT) with 561nm laser illumination after retro-orbital injection of Rose Bengal (Fig 1A). At 24 hours after the induction, a greatly increased level of reactive oxygen species (ROS) and cell death in the cerebral cortical area nearby the PT induction site were revealed by *in vivo* labeling of dihydroethidium (DHE) and propidium iodide (PI), respectively (Fig 1B and C). At 30 days after the PT, the cerebral cortex with the microinfarct was harvested and optically cleared for 3D volumetric imaging analysis. Interestingly, a strong second harmonic generation (SHG) signal indicating a formation of fibrous collagen-rich glial scar(Esquibel, Wendt et al., 2020, Perry, Burke et al., 2012) was detected by two-photon microscope (TPM) in the cerebral microinfarct as shown in 3D volume reconstructed images (Fig 1D). Cross-sectional Z-stack image sequence revealed accumulation of the collagen fiber at the periphery of the microinfarct (Appendix Fig S1). In addition, an almost complete loss of neuronal nuclei (NeuN) inside the microinfarct was confirmed with 3D volumetric imaging (Fig 1E and Movie EV1), indicating a focal neuronal loss associated with the microinfarct.

**Figure 1.**
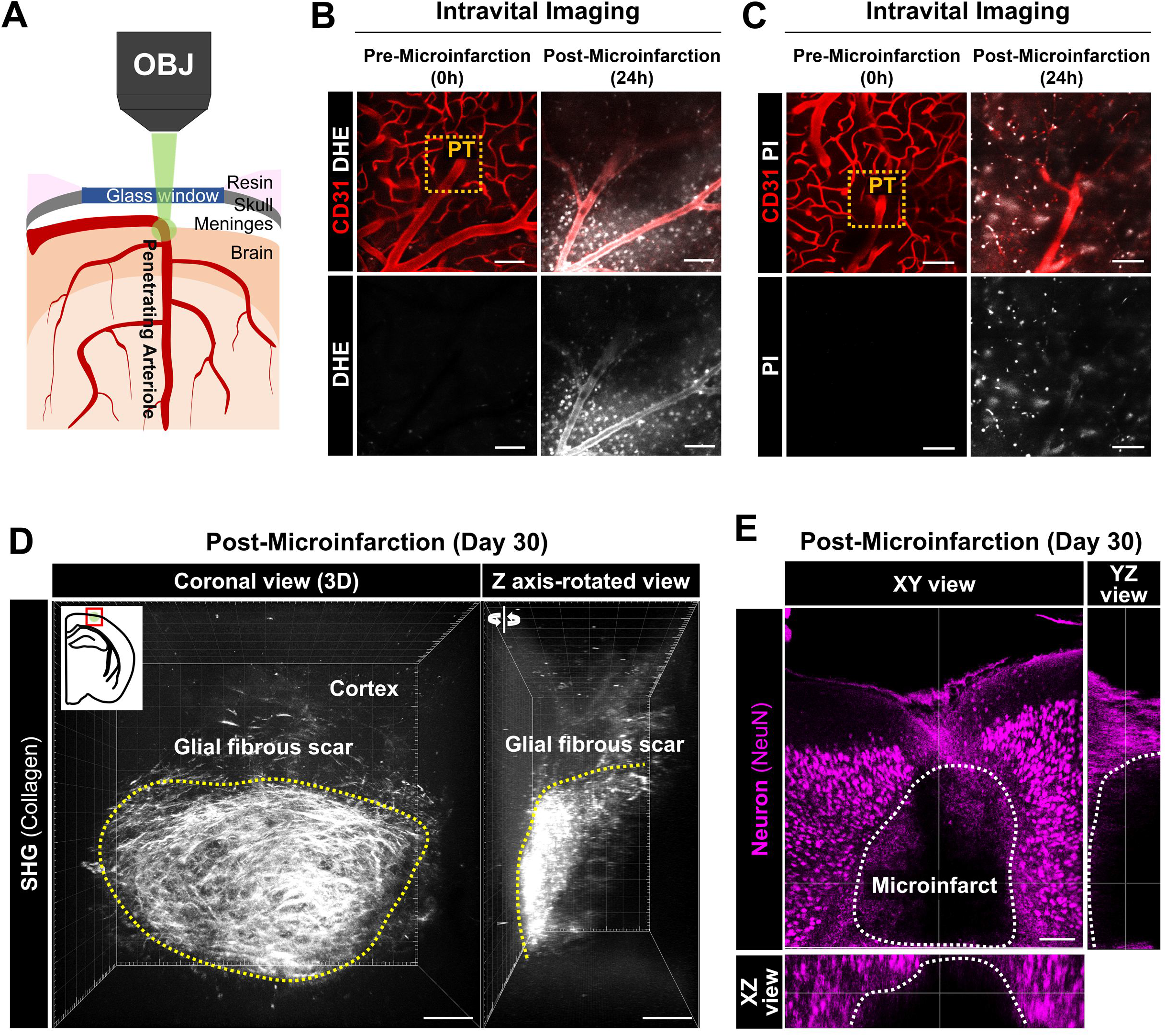
Cerebral microinfarction leads to glial scar formation in cerebral cortex. (**A**) Schematic of cerebral microinfarction model based on the selective photothrombosis (PT) in penetrating arteriole induced by focused laser beam through objective lens (OBJ). (**B** and **C**) Increased reactive oxygen species (detected by dihydroethidium; DHE, B) and cell death (detected by propidium iodide; PI, C) around the PT site (orange-dotted square). At 24 hrs after microinfarct induction, the PT site was identified based on vasculature labeled with CD31 antibody. (**D**) 3D reconstructed coronal-view and Z axis-rotated view images of optically cleared brain slice showing second harmonic generation (SHG) signal from collagen at 30 days after microinfarct induction. (**E**) XY/XZ/YZ view images of optically cleared brain slice showing the loss of neuronal nuclei (NeuN) in the microinfarct (white-dotted line) at 30 days after microinfarct induction. Scale bars = 100 μm.

### Focal atrophy with gradual collagen deposition and astrocyte loss in cerebral microinfarct

To investigate cellular-level alterations following microinfarction, we performed a longitudinal repetitive intravital imaging of the cerebral cortex for 30 days after the induction of PT in a live mouse implanted with the cranial imaging window. To simultaneously visualize astrocyte and pericyte composing the neurovascular unit (NVU), we used a double transgenic mouse by cross-breeding a pericyte-reporter NG2-DsRed mouse and an astrocyte-reporter Aldh1l1-GFP mouse (Appendix Fig S2). In the sham control group (n=5), no identifiable cellular-level alteration was observed up to 30 days in the same location of cerebral cortex (Appendix Fig S3). In the microinfarction group (n=5), acute tissue expansion followed by gradual tissue shrinkage was observed (Fig 2A, upper panel). A vasculature map identified on the bases of arteriole and venule identified by NG2-DsRed and CD31 labeling (Fig 2A, lower panel) showed that an acute ∼1.5 folds expansion of the brain tissue, indicating a severe edema, for 1-2 days followed by a gradual shrinkage to the baseline at 5-8 days and then a significant 50% reduction of the brain tissue, indicating a focal brain atrophy, for 2-4 weeks (Fig 2B). Next, to identify the time-courses of fibrous glial scar formation and astrocyte changes, we imaged optically cleared 500 μm thick brain sections obtained from Aldh1l1-GFP mice after the microinfarct induction. Aldh1l1^+^ astrocytes almost completely disappeared inside the microinfarct at day 5 and failed to repopulate at day 30 (Fig 2C, upper panel). At day 5 with the astrocyte loss, SHG signal started to be detected in the microinfarct where no Aldh1l1^+^ astrocyte was observed, which progressively increased at day 11, 20 and 30 (Fig 2C and D).

**Figure 2.**
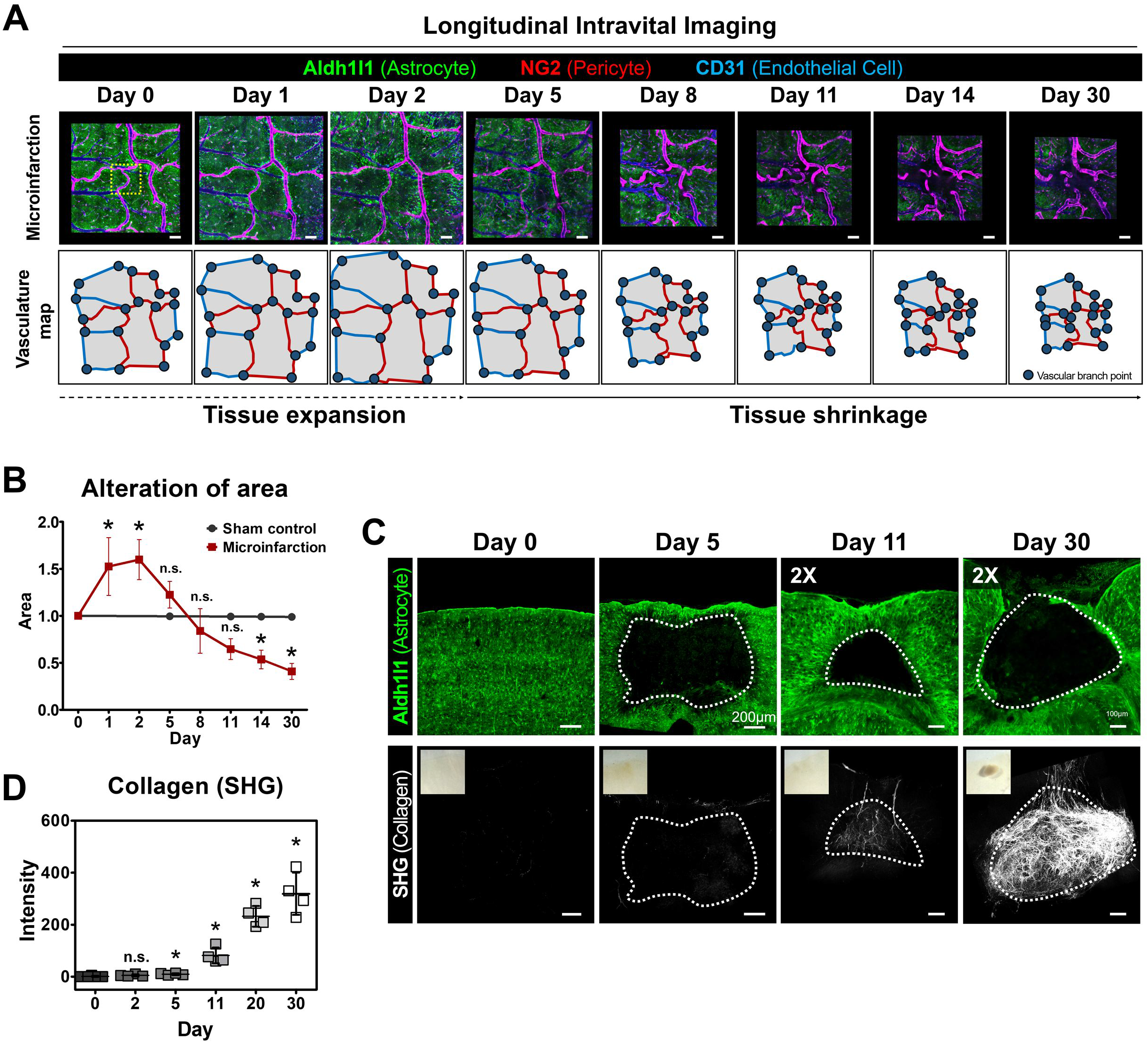
Longitudinal imaging reveals a focal atrophy with gradual collagen deposition and an astrocyte loss after the microinfarct induction. (**A**) Representative longitudinal intravital images after the microinfarct induction with photothrombosis (PT) for 30 days and vasculature map identified on the bases of venules and arterioles (green, Aldh1l1-GFP; red, NG2-DsRed; blue, CD31). (**B**) Tissue area alteration (n = 5 mice per each group, *\*P* < 0.05). (**C**) Representative maximal intensity projection mosaic images of brain section showing the loss of Aldh1l1^+^ astrocytes (green) and fibrous collagen deposition (white; SHG) in the microinfarct. (**D**) Fibrous collagen (SHG) in the microinfarct (n = 4 mice per each day, *\*P* < 0.05). Graphs are presented as mean ± s.d. Scale bars = 100μm.

### Acute loss of astrocyte and neuron in the microinfarct

At 2 days after the induction of photothrombosis, we identified that Aldh1l1^+^ astrocytes and their processes disappeared from the microinfarct lesion (Fig 3A). To observe the cellular-level dynamics of the astrocyte loss, a time-lapse intravital imaging of Aldh1l1^+^ astrocyte and CD31^+^ EC was performed for 12 hours starting from 1 day after the microinfarct induction, which revealed a sequential disappearance of juxtavascular Aldh1l1^+^ astrocytes along the blood vessel (Fig 3B and Movie EV2). In the sham control group, no alteration of Aldh1l1^+^ astrocyte was observed up to 30 days in the same location of cerebral cortex (Appendix Fig S4). In addition, the distribution of astrocyte and neuron in and around the microinfarct were identified by immunostaining with NeuN and glutamate transporter 1 (GLT-1), respectively. At day 0, both GLT-1^+^ astrocyte and NeuN^+^ neurons uniformly distributed in cortex (Fig 3C and Appendix Fig S5). At day 2, the DAPI^+^ nuclei number in the microinfarct significantly decreased by half (Fig 3D). Astrocytes in the microinfarct were poorly labelled by GLT-1 as compared to those in the peri-infarct whereas those in the normal area had similar level of staining to day 0. In addition, the number and the size of nuclei in NeuN^+^ neurons decreased greatly (Fig 3E and F), suggesting that these neurons were in the process of death.

**Figure 3.**
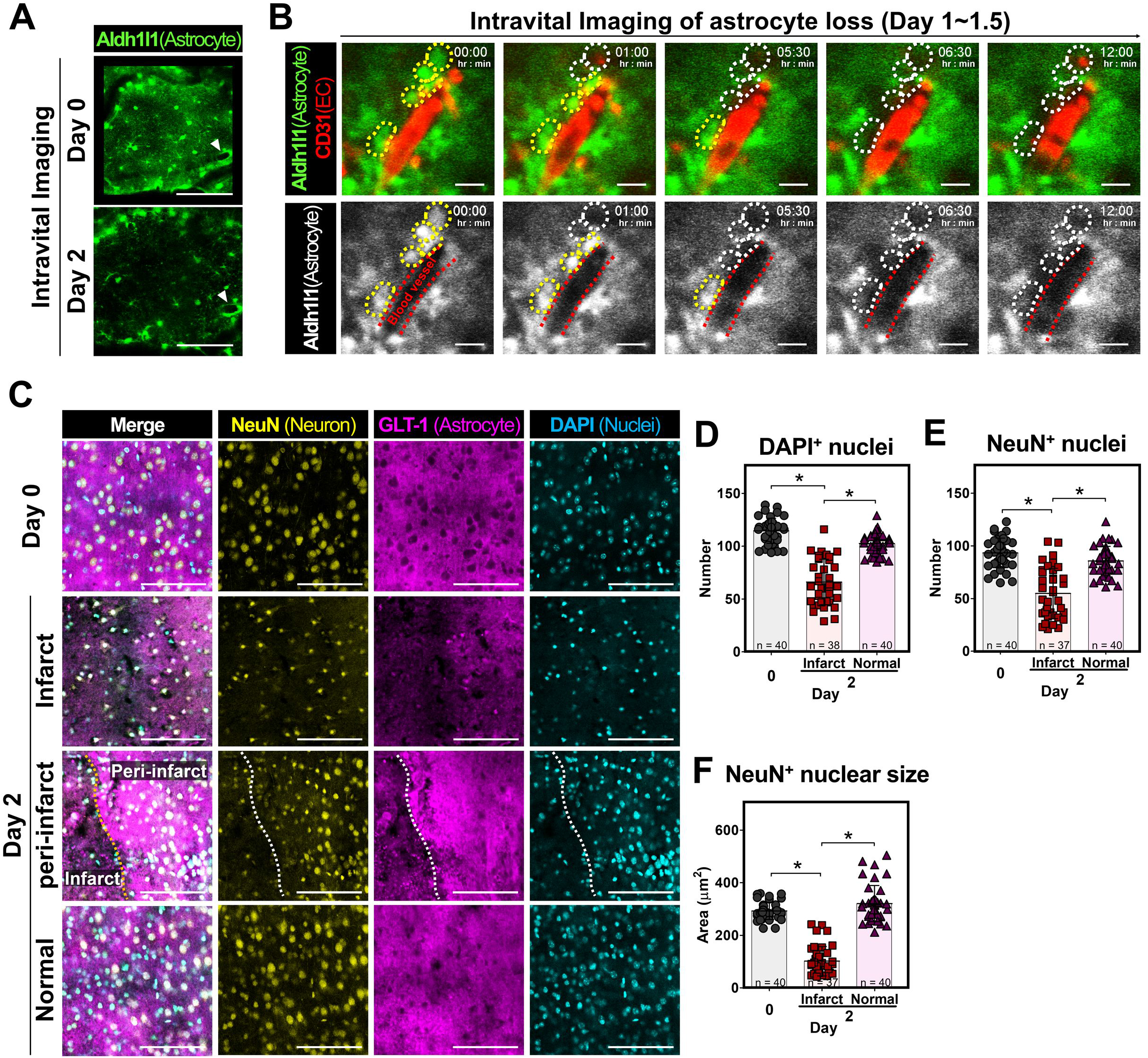
Acute loss of astrocyte and neuron after the microinfarct induction. (**A**) Representative intravital images showing acute loss of Aldh1l1^+^ astrocyte at 2 days after the microinfarct induction (green, Aldh1l1-GFP). (**B**) Twelve-hours time-lapse intravital images showing the disappearance of Aldh1l1^+^ astrocytes at 1∼1.5 day after the microinfarct induction (green, Aldh1l1-GFP). Yellow-dotted and white-dotted lines marked the astrocytes before and after disappearance, respectively. Red-dotted line marked the blood vessel (CD31). (**C**) Representative images of NeuN and glutamate transporter 1 (GLT-1) expression in the microinfarct, peri-infarct and normal area in brain at day 0 and day 2 (yellow, NeuN; magenta, GLT-1; cyan, DAPI). (**D** and **E**) The number of DAPI^+^ nuclei (**D**) and NeuN ^+^ nuclei (**E**) (n > 38 spots in 3 mice; \**P* < 0.05). (**F**) The size of NeuN ^+^ nuclei (n > 37 spots in 3 mice; \**P* < 0.05). Graphs are presented as mean ± s.d. Scale bars = (**A** and **C**) 100μm, (**B**) 10μm.

### Acute loss of pericyte coverage and recovery

To examine how pericyte responded following a microinfarction, a longitudinal intravital imaging of NG2^+^ pericyte at same cortex area in a single mouse was performed for 11 days after the microinfarct induction. An acute decrease in the pericyte coverage over CD31^+^ blood vessels was observed at day 2, which was at least partially recovered at day 5 (Fig 4A). In the sham control group, no changes in pericyte processes length or vascular coverage was observed up to 30 days (Fig 4B and Appendix Fig S6). In the microinfarct, the processes length of NG2^+^ pericytes significantly decreased by more than 48% from day 1 to day 3 after the microinfarct induction then recovered at day 5 and maintained until day 11 (Fig 4B). To observe the cellular-level dynamics in the recovery of pericyte coverage, a time-lapse intravital imaging of NG2^+^ pericyte expressing histone H2B-GFP in their nuclei was performed for 24 hours starting from 3.5 days after the microinfarct induction. Proliferation of juxtavascular NG2^+^ pericyte (Fig 4C and Movie EV3) and processes elongation along the vessel (Fig 4D and Movie EV4) were clearly imaged, suggesting that both processes contributed to the recovery of pericyte coverage from day 3 to day 5 after the microinfarct induction.

**Figure 4.**
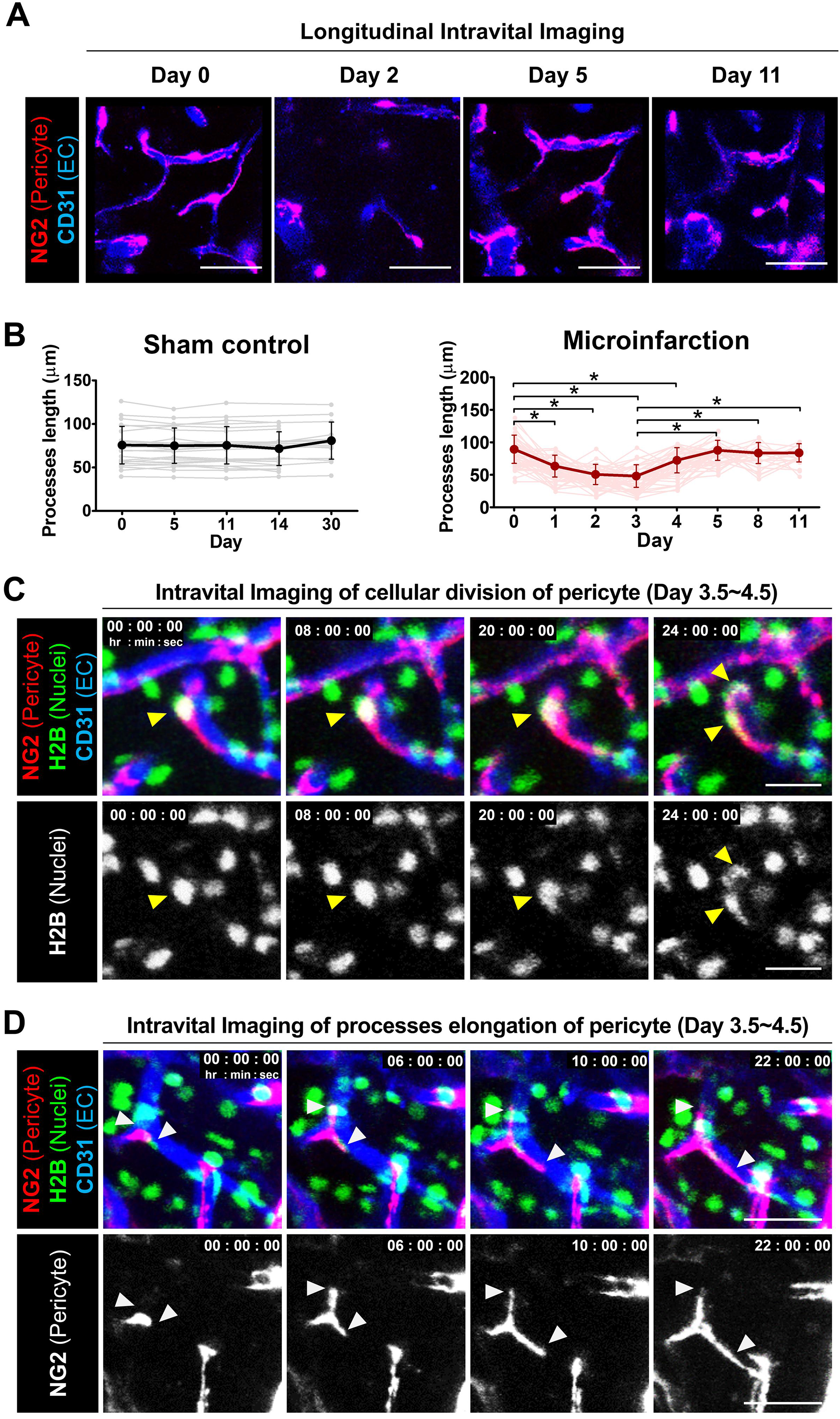
Acute loss of pericyte coverage after the microinfarct induction and recovery of coverage through cellular division and processes elongation of pericyte. (**A**) Representative longitudinal intravital images of NG2^+^ pericytes and ECs for 11 days (red, NG2-DsRed; blue, CD31). (**B**) NG2^+^ pericyte processes lengths (n > 25 pericyte in 5 mice, \**P* < 0.05). (**C** and **D**) Twenty-four-hours time-lapse intravital images of NG2^+^ pericytes expressing H2B-GFP in nuclei showing cellular divisions marked by yellow arrowheads (**C**) and processes elongation marked by white arrowheads (**D**) at 3.5∼4.5 days after the microinfarct induction (green, H2B-GFP; red, NG2-DsRed; blue, CD31). Graphs are presented as mean ± s.d. Scale bars = 50μm.

### Impairment of vascular flow and leakage followed by partial recovery

To examine an impairment in cerebral vascular perfusion and leakage following a microinfarction, a longitudinal intravital imaging of blood flow at same cortex area in a single mouse was performed for 5 days in the peri-infarct after the microinfarct induction (Fig 5A). The blood flow was visualized by intravenously injected 2MDa FITC-dextran and 3kDa TRITC-dextran right before each imaging experiment at days 0, 3 and 5. At day 3, a leakage of 3kDa TRITC-dextran but not 2MDa FTIC-dextran from blood vessel to brain parenchyma was observed, suggesting a compromised blood-brain-barrier (BBB) integrity at least for smaller-sized molecules in the peri-infarct causing the edema observed at same time point (Fig 2A). Interestingly, the vascular leakage of 3kDa TRITC-dextran was not observed at day 5. The fluorescence intensity of 3kDa TRITC-dextran detected in brain parenchyma showed similar changes, significant increase at day 3 followed by decrease to the baseline at day 5, while the fluorescence intensity of 2MDa FTIC-dextran showed no changed from day 0 to day 5 (Fig 5B and C). In addition, an impairment and recovery of vascular flow in cerebral microvasculature at peri-infarct area next to the site of photothrombosis was observed by a longitudinal intravital imaging for 5 days after the microinfarct induction. The blood flow and vasculature were visualized by intravenously injected Alexa Flour 647-dextran and a photo-convertible protein ‘Kaede’(Tomura, Yoshida et al., 2008) with NG2-DsRed^+^ pericyte, respectively. For individual capillary branches followed for 5 days, four distinguishable dynamic patterns of changes in blood flow and vasculature were identified, including maintained flow, recovered flow, blocked flow, and vessel regression (Fig 5D and Appendix Fig S7). Among a total of 157 capillary branches identified at day 0 with blood flow (mouse n=4), 83 capillary branches were observed to have no blood flow, categorized as blocked flow, at day 2. In each mouse, the proportion of individual capillary branches maintaining blood flow at day 2 ranged between 30.5 % to 70 % (Fig 5E). Interestingly, 18.9% of capillary branches observed with blood flow at day 2 were found to have no blood flow or have disappeared at day 5, which were categorized as blocked flow and vessel regression, respectively. On the other hand, 61.4% of capillary branches observed with no blood flow at day 2 were found to recover blood flow at day 5, which was categorized as recovered flow. These combined changes in the blood flow in capillary branches resulted in partial recovery of blood flow, from 47.6% at day 2 to 74% at day 5 (Fig 5E).

**Figure 5.**
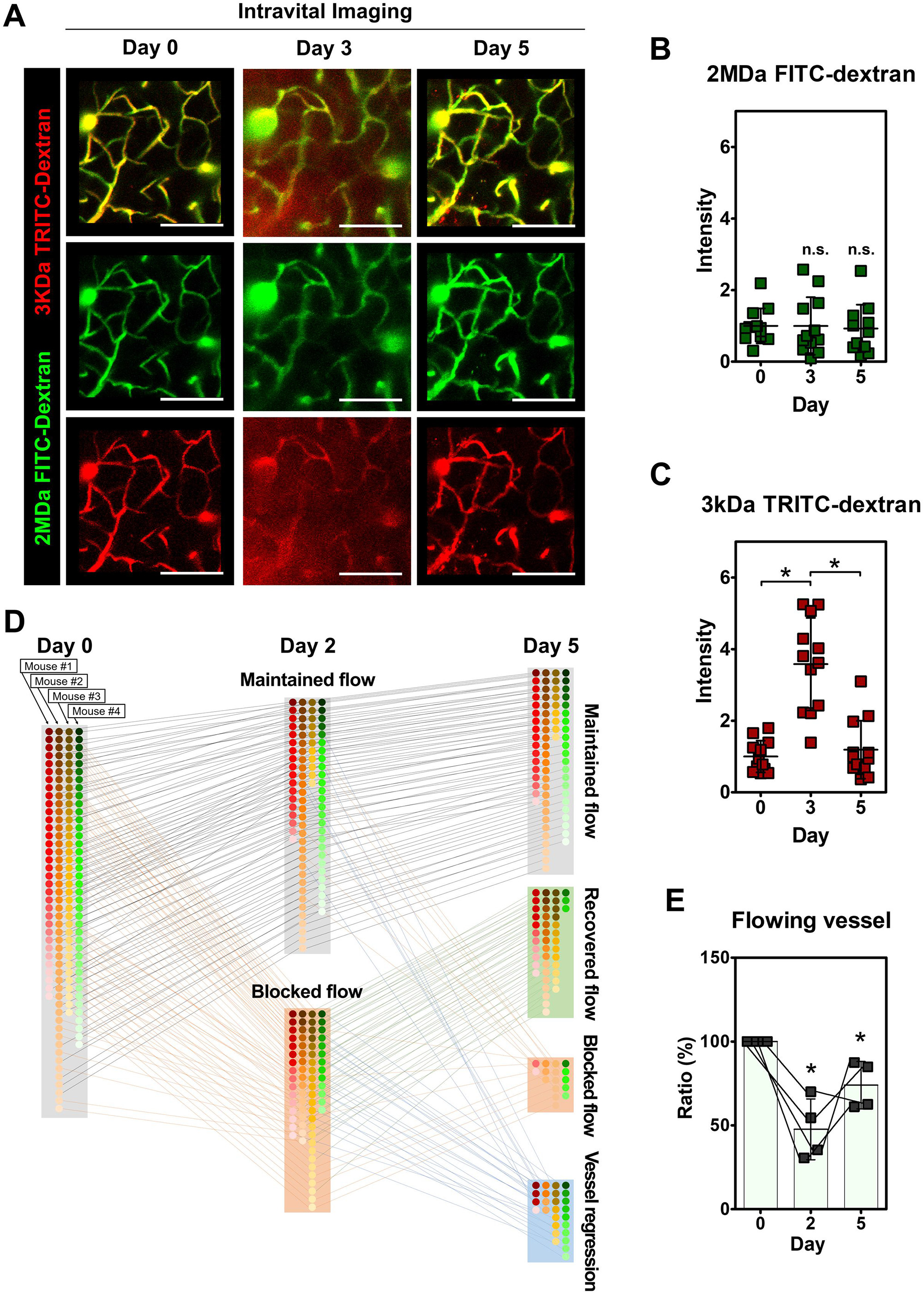
Acute impairment and partial recovery of vascular flow and leakage after the microinfarct induction. (**A**) Representative longitudinal intravital images of vascular flow and leakage for 5 days after the microinfarct induction (green, 2MDa FITC-dextran; red, 3kDa TRITC-dextran). (**B** and **C**) Fluorescence intensity of 2MDa FITC-dextran (**B**) and 3kDa TRITC-dextran (**C**) in cortex parenchyma of (A) (n = 12 spots in 3 mice; \**P* < 0.05). (**D**) Longitudinal tracking analysis of individual capillary branches for 5 days after the microinfarct induction (n = 83 branches in 4 mice). (**E**) Ratio of capillary branches with blood flow (\**P* < 0.05). Graphs are presented as mean ± s.d. Scale bars = 100μm.

### Accumulation of GFAP^+^ reactive astrocytes at peri-infarct

At day 4 after the microinfarct induction, a ∼1.5 fold increase in the size of astrocyte soma was observed at peri-infarct area (Fig 6A and B). To identify the increase of astrocyte reactivity, an immunostaining of glial fibrillary acidic protein (GFAP) and SHG imaging were performed at day 30 after the microinfarct induction (Fig 6C and D). Based on the loss of Aldh1l1^+^ astrocytes and the formation of fibrous collagen-rich glial scar, the infarct area was delineated to identify the peri-infarct area (Fig 6C). Remarkably, GFAP^+^ reactive astrocytes appeared to be highly accumulated at the peri-infarct area up to ∼200 μm away from the infarct border (Fig 6D). Serial observation of the peri-infarct area revealed that GFAP^+^ reactive astrocytes started to appear at the peri-infarct area at 2 days after the microinfarct induction and remained until 30 days with the glial scar formation (Fig 6E and F). In addition, Aldh1l1^+^ astrocyte in the peri-infarct area significantly increased from day 5 and remained until 30 days (Fig 6G and H). Sholl analysis of individual Aldh1l1^+^ astrocytes accumulated at the peri-infarct showed a significant increase of process lengths and a decrease of process number (Appendix Fig S8), implying an increased astrocyte reactivity accompanying the increased GFAP. Furthermore, an immunostaining of lipocalin2 (LCN2) showed an accumulation of LCN2^+^ GFAP^+^ astrocytes, presumably severe reactive astrocytes (Bi, Huang et al., 2013), co-localized with Aldh1l1^+^ astrocytes at the peri-infarct (Fig 6I).

**Figure 6.**
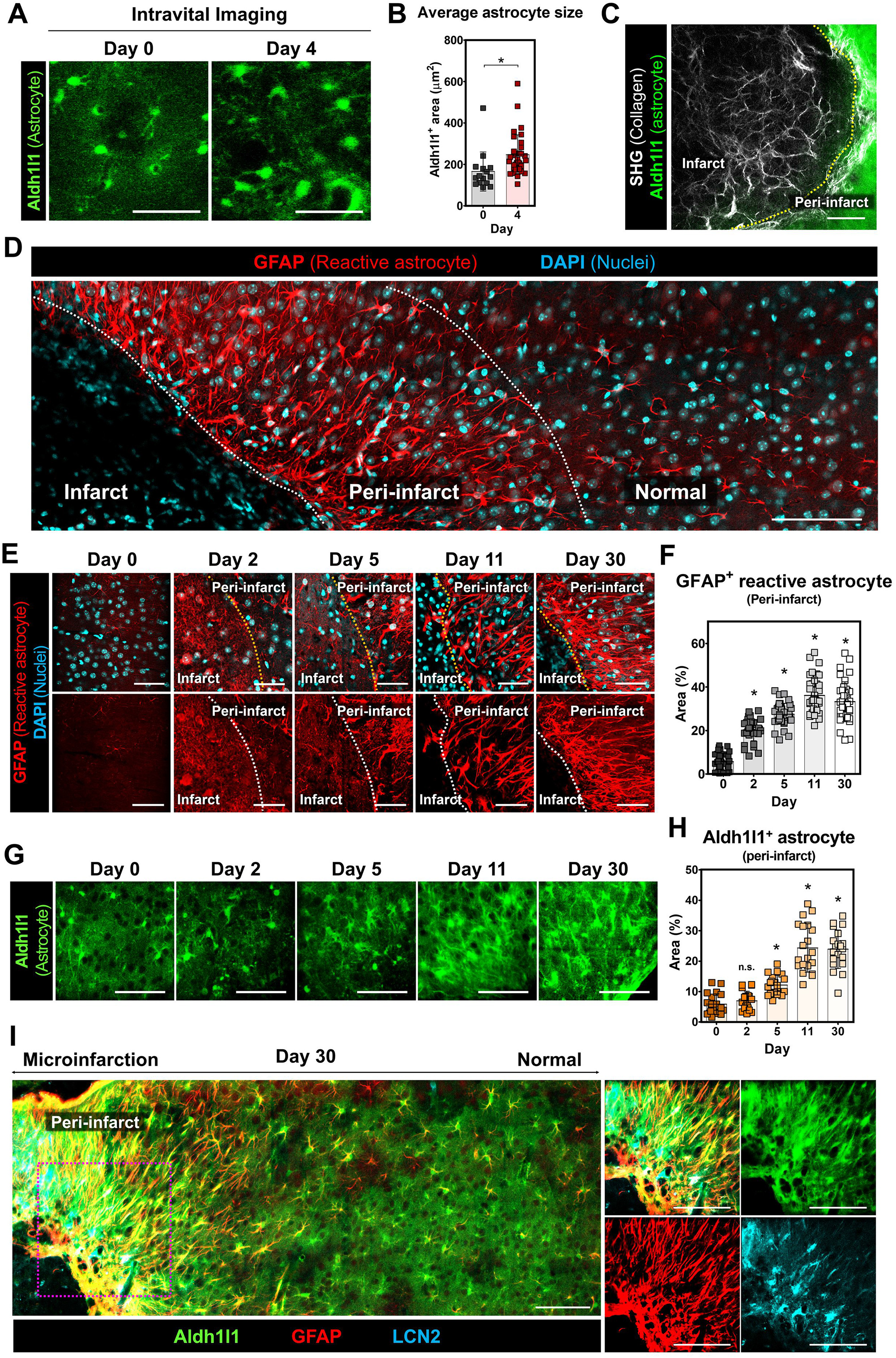
Accumulation of GFAP^+^ reactive astrocytes at peri-infarct area. (**A**) Representative intravital images of Aldh1l1^+^ astrocyte with enlarged soma after the microinfarct induction (green, Aldh1l1-GFP). (**B**) Size of Aldh1l1^+^ astrocyte (day 0, n = 15 cells; day 4, n = 37 cells; *\*P* < 0.05). (**C)** Fibrous collagen (detected by SHG) deposition and loss of Aldh1l1^+^ astrocyte in the microinfarct at 30 days after the microinfarct induction. (**D**) Representative wide-area mosaic image showing accumulation of GFAP^+^ reactive astrocytes at the peri-infarct area at 30 days after the microinfarct induction (red, GFAP; cyan, DAPI). (**E**) Serial images of accumulation of GFAP^+^ reactive astrocytes at the peri-infarct area for 30 days after the microinfarct induction. (**F**) GFAP^+^ reactive astrocyte area at the peri-infarct (n > 39 spots in 4 mice; *\*P* < 0.05). (**G**) Serial images of Aldh1l1^+^ astrocytes at the peri-infarct for 30 days after the microinfarct induction (green, Aldh1l1-GFP). (**H**) Aldh1l1^+^ astrocyte area at the peri-infarct (n = 20 spots in 2 mice; \**P* < 0.05). (**I**) Representative wide-area mosaic and magnified images of GFAP^+^LCN2^+^ severe reactive astrocytes at day 30 after the microinfarct induction (green, Aldh1l1-GFP; red, GFAP; blue, LCN2). Scale bars = (**A** and **E**) 50μm, (**C**,**D**,**G** and **I**) 100μm.

### Reduced fibrous glial scar formation through an oral administration of KDS2010

To investigate the effect of an intervention on the astrocyte reactivity in the fibrous glial scar formation, KDS2010, a monoamine oxidase B (MAO-B) inhibitor in reactive astrocyte, was orally administered with an oral zonde needle (10 mg/kg per day) for 30 days after the microinfarct induction. While the size of infarct with astrocyte and neuronal loss was not decreased, a significant ∼ 0.4 fold decrease in the accumulated collagen was observed (Fig 7A-C). Additionally, Sholl analysis of individual Aldh1l1^+^ astrocytes accumulated at the peri-infarct showed a significant decrease of process lengths and an increase of process number (Fig 7D-F), implying a decreased astrocyte reactivity. Concurrently, LCN2^+^ GFAP^+^ severe reactive astrocytes decreased at the peri-infarct area, while the total GFAP^+^ astrocytes accumulation did not change (Appendix Fig S9).

**Figure 7.**
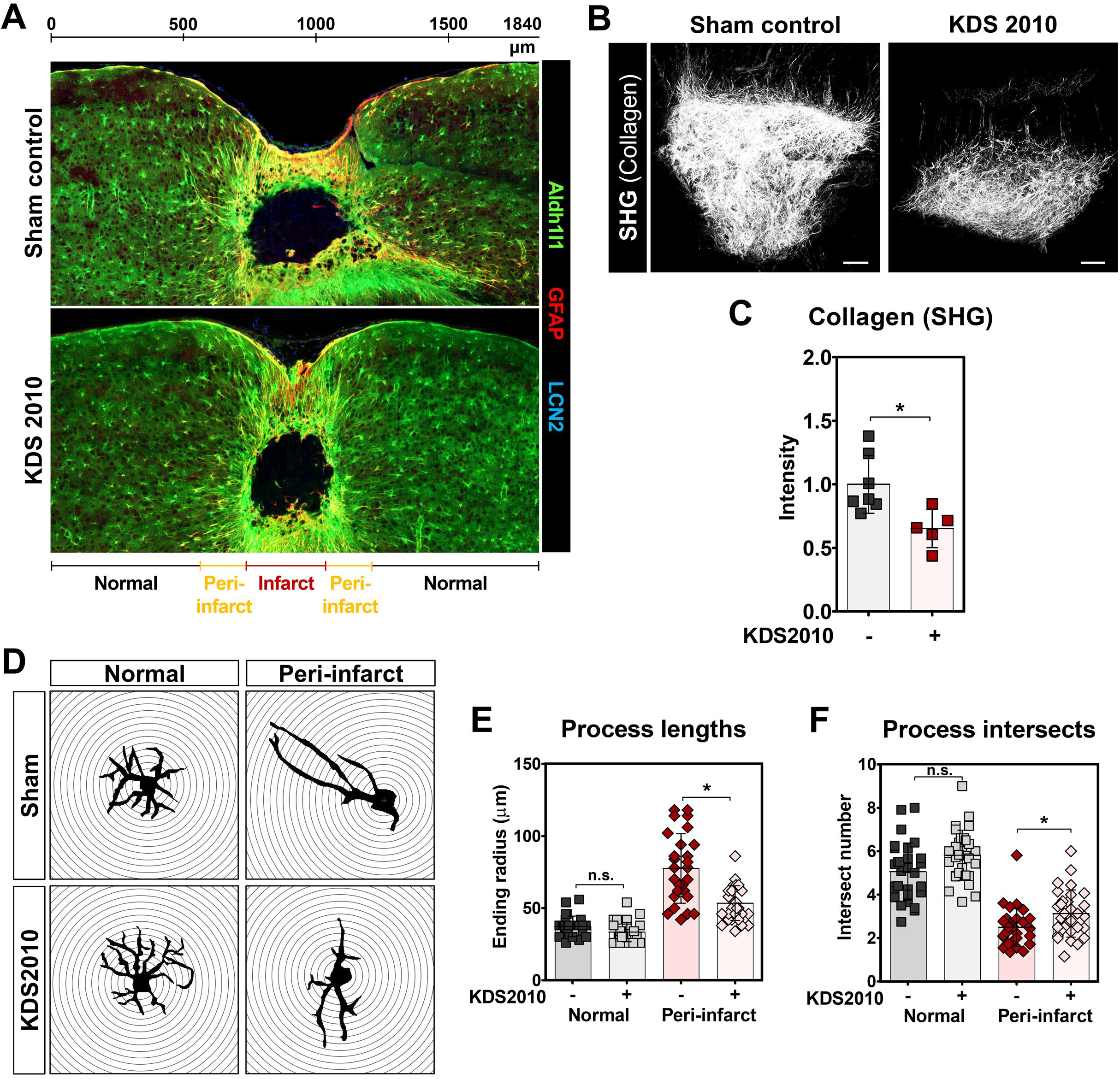
Reduced glial scar formation through an oral administration of KDS2010. (**A**) Representative wide-area mosaic images of brain section with microinfarct from non-treated sham control group and KDS2010-treated group at 30 days after the microinfarct induction. (**B**) Representative maximal intensity projection images of fibrous collagen in the microinfarct. (**C**) Quantification of fibrous collagen, measured as intensity of SHG, in the microinfarct (sham control, n = 7 mice; KDS2010-treated, n = 5 mice; \**P* < 0.05). (**D**) Representative results of Sholl analysis of Aldh1l1^+^ astrocytes in normal and peri-infarct area of sham control and KDS2010-treated group. (**E** and **F**) Process lengths (**E**) and process intersects (**F**) of individual astrocytes (n= 30, \**P* < 0.05). Graphs are presented as mean ± s.d. Scale bars = 100μm.

## Discussion

Conventional detection methods for cerebral microinfarcts are neuropathological examination, diffusion-weighted imaging (DWI), and high-resolution structural MRI(van Veluw et al., 2017). Neuropathological examination can detect the smallest hyperacute to chronic microinfarcts in a brain autopsy(Westover, Bianchi et al., 2013), DWI can detect a sub-millimeter sized small microinfarcts(Keir & Wardlaw, 2000), and high-resolution structural MRI using 7T(van Veluw, Jolink et al., 2014) and 3T MRI(Hilal, Sikking et al., 2016, van Veluw, Hilal et al., 2015) can detect an 1-2 mm sized microinfarcts in the cortical grey matter of human brain. Notably, a post-mortem histological examination of brain sections of individuals with dementia showed a neuronal loss, collagen 4-positive microvessel and GFAP-positive astrocyte in the microinfarct core detectable by T2-weighted imaging(Yilmazer-Hanke, Mayer et al., 2020). Histological examination of dissected brain tissue has the limitation in analyzing spatiotemporal changes in the microinfarct *in vivo* as it only provides information at the end-point. While non-invasive DWI and MRI can follow the microinfarct *in vivo* over long periods of time, the detectable sizes of microinfarct are limited to larger than ∼ 1mm and the follow-up time-points are relatively limited due to the high cost. Although its application is currently limited to animal model, the intravital microscopy capable of an *in vivo* longitudinal observation of brain based on chronic imaging window(Holtmaat, Bonhoeffer et al., 2009) can be a better method to mechanistically investigate the spatiotemporal changes after the onset of small microinfarct in sub-cellular resolution *in vivo*. To note, intravital longitudinal imaging of mouse cerebral cortex enabled by the chronic imaging window has revealed a myelin degeneration and internodes loss in aged mouse(Hill, Li et al., 2018), an impairment of short-term memory function with reduction of blood flow in Alzheimer’s disease (AD) models APP/PS1 and 5xFAD mice(Cruz Hernandez, Bracko et al., 2019), and neurovascular uncoupling, resulting from pericyte deficiency in mice with a loss-of-function mutation of platelet-derived growth factor receptor-β (*Pdgfrb*^+/−^)(Kisler, Nelson et al., 2017).

In this study, we successfully performed a longitudinal imaging of mice for 1 month after the induction of microinfarction with a custom-built intravital confocal and two-photon microscope. The longitudinal imaging revealed an acute tissue expansion with an acute astrocyte loss at day 2 followed by gradual tissue shrinkage for 1 month, resulting in a cerebral atrophy with a neuronal loss and a persisted astrocyte loss in the microinfarct (Fig 2 and 3). Interestingly, a fibrous collagen-rich glial scar formation was observed in the microinfarct with second harmonic generation (SHG) imaging using the two-photon microscope (Fig 1D). SHG is a nonlinear optical process that two photons are combined to form a new photon with twice the energy, which efficiently occurs in noncentrosymmetric molecules such as the collagen in biological tissue enabling a visualization of collagen without additional labeling(Mostaco-Guidolin, Rosin et al., 2017, Zipfel, Williams et al., 2003). A label-free SHG imaging have been widely used to monitor the fibrillar collagen in specimens of various fibrotic diseases in skin(Chen, Chen et al., 2009), liver(Goh, Leow et al., 2019), lung(Kottmann, Sharp et al., 2015), carotid artery(Megens, oude Egbrink et al., 2008), eye(Tan, Sun et al., 2007), tendon and ligaments(Doras, Taupier et al., 2011). In brain tissue, SHG imaging was used to investigate the structure and polarity of microtubules in neuronal processes(Dombeck, Kasischke et al., 2003, Van Steenbergen, Boesmans et al., 2019), and collagen deposition in glioblastoma(Jiang, Wang et al., 2017). However, to the best of our knowledge, neither the fibrous collagen accumulation and scar formation associated with the brain microinfarcts nor the SHG imaging of them has been reported.

In the central nervous system (CNS), fibrotic scar has been reported to be generated by scar-forming cells such as EC, immune cells, fibroblasts, and astrocyte in focal damaged tissue associated with neurodegenerative diseases(D’Ambrosi & Apolloni, 2020, Dorrier, Aran et al., 2021), spinal cord injury(Hara, Kobayakawa et al., 2017) and brain injuries(Frik, Merl-Pham et al., 2018, Huang, Wu et al., 2014). After CNS injury, fibrotic scar segregated the damaged lesion from normal tissue to prevent the spread of cellular damage, however it also impaired axonal regeneration and thus limited functional recovery(Kawano, Kimura-Kuroda et al., 2012). Reducing fibrotic scar formation by ablating CNS fibroblasts reduced motor disability in experimental autoimmune encephalomyelitis (EAE) mouse model of multiple sclerosis(Dorrier et al., 2021). Moreover, suppression of scar formation by intracerebral injection of 2,2’-dipyridyl (DPY) to inhibit the type VI collagen formation promoted axonal regeneration after traumatic brain injury(Yoshioka, Hisanaga et al., 2010). These reports suggest that elucidating the underlying cellular and molecular mechanisms in fibrous glial scar formation surrounding the microinfarct could potentially led to the development of therapeutics to alleviate an adverse impact of the microinfarct.

A significant increase of astrocyte reactivity can be induced after CNS insults, such as trauma, stroke, infection, and neurodegenerative diseases(Sofroniew, 2009). In this process, astrocytes undergo dramatic changes in their morphology, hypertrophy of soma and processes, and in molecular expression including elevated glial fibrillary acidic protein (GFAP), and then they can induce a glial scar formation in CNS in severe cases(Hara et al., 2017, Sofroniew, 2009, Wanner, Anderson et al., 2013). In our study, at 4 days after the microinfarct induction, a ∼1.5 fold increase in the size of astrocyte soma was observed at peri-infarct area (Fig 6A and B), suggesting a reactive astrocyte change. Consistently, at the peri-infarct area, GFAP^+^ reactive astrocytes started to be observed from 2 days after the microinfarct induction and were highly accumulated until 30 days with fibrous collagen accumulation (Fig 6C-F). Furthermore, Lipocalin 2 (LCN2), a maker of severe reactive astrocyte, was co-expressed in GFAP^+^ reactive astrocyte at the peri-infarct area (Fig 6I). LCN2 has been reported as an autocrine factor for reactive astrocytosis(Lee, Park et al., 2009), has a neurotoxicity, and can be secreted from GFAP^+^ reactive astrocytes in neurodegenerative disease animal model(Bi et al., 2013). Additionally, severe reactive astrocytes recognized by co-expression of LCN2 and GFAP were characterized as scar-forming reactive astrocytes in spinal cord injury animal model(An, Lee et al., 2020).

Amelioration of astrocyte reactivity by inhibiting the Janus kinase 2 (JAK2)-STAT3 pathway can elicit a beneficial effect in spatial learning in an Alzheimer’s disease (AD) model(Ceyzeriat, Ben Haim et al., 2018). Removal of hydrogen peroxide (H_2_O_2_) in severe reactive astrocyte by AAD-2004, a potent radical scavenger, can prevent spatial memory impairment in an AD model caused by severe reactive astrocytes(Chun, Im et al., 2020). KDS2010, a newly developed highly-selective reversible MAO-B inhibitor for reactive astrocytes, has demonstrated a potent therapeutic benefit in alleviating learning and memory impairments in APP/PS1 AD mice by attenuating the astrocytic reactivity(Park, Ju et al., 2019). In our study, the administration of KDS2010 significantly reduced the collagen deposition in the microinfarct by half (Fig 7B and C). Consistently, KDS2010 could decrease the reactivity of astrocyte by reducing LCN2 expression in GFAP^+^ reactive astrocytes (Fig 7D and E) and process lengths of individual Aldh1l1^+^ astrocyte and by increasing the process intersects of Aldh1l1^+^ astrocyte (Appendix Fig S8). However, unfortunately, neither a reduction in the size of microinfarct lesion nor a recovery of astrocyte loss in the microinfarct was achieved (Fig 7A) and a noticeable amount of collagen still remained in the microinfarct (Fig 7B and C). These limited improvements in pathological features with KDS2010 treatment could be attributed to the infiltrated immune cells such as CX3CR1^+^ immune cells reported to be highly accumulated in the microinfarct after photothrombotic occlusion(Lubart, Benbenishty et al., 2021) or myelin debris engulfment of endothelial cells(Zhou, Zheng et al., 2019) and proliferative CNS fibroblasts(Dorrier et al., 2021). Additional intervention strategies targeting other scar-forming factors would be needed to further decrease the fibrous scar formation after the onset of microinfarct and improve pathological features. On the other hand, brain stimulation could be considered as an adjuvant therapy. Optogenetic neuronal stimulation in the lateral cerebellar nucleus showed a persistent recovery on the rotating beam test for motor/sensory function after stroke(Shah, Ishizaka et al., 2017). Electrical vagus nerve stimulation was reported to reduce infarct volume and improve a neuronal function after cerebral ischemic injury(Xiang, Wang et al., 2015).

In human, cortical microinfarcts were consistently observed in the proximity of amyloid angiopathy in AD patient(Okamoto, Ihara et al., 2009). AD patients have more microinfarcts revealed by 7-T magnetic resonance (MR) imaging(van Rooden, Goos et al., 2014). Yet, the association between cerebral microinfarction and AD is still unclear. Interestingly, we found reactive astrocytes identified by hypertrophy around small amyloid plaque located at capillary vessel with blocked blood flow in the brain of 5.5 months old APP/PS1 AD mouse cross-bred with Aldh1l1-GFP x NG2-DsRed mouse by intravital imaging with the chronic cranial imaging window (Appendix Fig S10A). Furthermore, the loss of pericyte coverage and astrocyte endfeet were observed at the same capillary vessel with the blocked blood flow. Notably, a repeated intravital imaging of the same capillary vessel after 1 month revealed that the blood flow was remained to be blocked and the losses of pericyte coverage and astrocyte endfeet persisted with the amyloid plaque (Appendix Fig S10B). Normally, 5.5 ∼ 6.5 months old APP/PS1 AD mice started to show non-detectable or mild symptoms of cognitive impairment (Webster, Bachstetter et al., 2014). Yet, at those periods, our imaging result showed that amyloid plaque could be spatially associated with impaired microcirculation, astrocyte reactivity, and NVU disruption, which could potentially play a role in the development of cerebral microinfarctions in AD patient(Morrone, Bishay et al., 2020, Okamoto et al., 2012, Suter, Sunthorn et al., 2002).

In this study, we longitudinally observe a complex cellular-level changes of astrocyte, pericyte and vascular functions and integrity after microinfarct induction. While acute loss of pericyte coverage, impaired vascular flow and integrity are partially recovered at day 5, the acute loss of astrocyte and neuron persists in the microinfarct at day 30. Notably, a fibrous collagen-rich glial scar formation is observed in the microinfarct with accumulation of GFAP^+^LCN2^+^ reactive astrocytes in the peri-infarct. Administration of KDS2010, a reversible inhibitor of monoamine oxidase B (MAO-B) in reactive astrocytes, significantly decreases the glial scar formation and astrocyte reactivity. Collectively, our longitudinal intravital imaging study of neurovascular pathophysiology in cerebral microinfarction demonstrate a susceptibility of astrocytes to ischemic brain injury and their reactive response leads to a long-term glial scar formation in cerebral cortex, which can be alleviated by suppressing the astrocyte reactivity.

## Methods

### Mice

All mice experiments were performed in accordance with the standard guidelines for the care and use of laboratory animals and were approved by the Institutional Animal Care and Use Committee (IACUC) of KAIST (approval No. KA2018-66). Mice used in this study were maintained in a specific pathogen-free facility of KAIST Laboratory Animal Resource Center. All mice were individually housed in ventilated and temperature & humidity-controlled cages (22.5 °C, 52.5 %) under 12-12 hours light-dark cycle and provided with standard diet and water ad libitum. For experimental use, 8–12 weeks old male mice (20∼30 g) were utilized in this study. C57BL/6 mice purchased from OrientBio (Korea) were used in this study. NG2-DsRed mice (Stock No. 008241, Jackson Laboratory) where DsRed is expressed under pericytes-specific NG2 promotor were purchased from the Jackson Laboratory. Aldh1l1-GFP mice expressing Green Fluorescence Protein (GFP) in most of astrocytes were generously provided by Dr. W. Chung (Korea Advanced Institute of Science and Technology, Korea). Aldh1l1-GFP x NG2-DsRed double transgenic mouse was produced by crossbreeding NG2-DsRed mouse and Aldh1l1-GFP mouse. NG2-DsRed x Kaede double transgenic mouse was generated by crossbreeding NG2-DsRed mouse and Kaede mouse (generously provided by Dr. Tomura and Dr. Miwa at Kyoto University, Japan), which universally express a photo-convertible fluorescent protein ‘Kaede’ (Tomura et al., 2008), and we performed the photoconversion at day 2 by 405nm laser illumination after the microinfarct induction to identify the presence of intermediate non-flowing vessel. NG2-DsRed x H2B-GFP double transgenic mouse was generated by crossbreeding NG2-DsRed mouse and H2B-GFP mouse (Stock No. 006069, Jackson Laboratory), which express GFP with a histone H2B gene. Finally, Aldh1l1-GFP x NG2-DsRed x APP/PS1 mouse was generated by crossbreeding NG2-DsRed, Aldh1l1-GFP mouse and APP/PS1 (MMRRC Stock No. 34832, Jackson Laboratory), which express a chimeric mouse/human amyloid precursor protein and a mutant human presenilin 1 associated with early-onset AD.

### Imaging system

Custom-built video-rate laser-scanning confocal and two-photon microscope system previously implemented (Ahn, Kong et al., 2019, Ahn, Choe et al., 2017, Choe, Hwang et al., 2013, Lee, Kong et al., 2020, Park, Choe et al., 2018, Park, Kim et al., 2019, Seo, Hwang et al., 2015) was used. Four continuous-wave laser modules with output wavelengths at 405nm (OBIS 405, Coherent), 488 nm (MLD488, Cobolt), 561 nm (Jive, Cobolt) and 640 nm (MLD640, Cobolt) were used as an excitation sources for confocal microscope. A femtosecond pulse Ti:Sapphire laser (690∼1050 nm, Chameleon Vision-S, Coherent) was used as a two-photon excitation source. A video-rate two-dimensional Raster-pattern laser-scanning was achieved by using a rotating polygonal mirror (MC-5, Lincoln Laser) for fast X-axis scanning and a galvanometer scanner (6230H, Cambridge Technology) for slow Y-axis scanning. Multi-color confocal fluorescence signals were detected by four photomultiplier tubes (PMT; R9110, Hamamatsu) through bandpass filters (FF01-442/46, FF02-525/50, FF01-600/37, FF01-685/40, Semrock). Multi-color two-photon fluorescence signals and second harmonics generation (SHG) signals were detected by four photomultiplier tubes (PMT; R11540, Hamamatsu) through bandpass filters (FF01-445/20, FF01-525/45, FF01-593/46, FF02-675/67, Semrock). Output signal of PMTs were digitally acquired by using a frame grabber (Solios, Matrox) and a custom-written image acquisition software based on the Matrox Imaging Library (MIL10, Matrox) for a real-time image acquisition, display and recording.

### Cranial imaging window preparation

Chronic cranial imaging window implantation (Holtmaat et al., 2009, Lee et al., 2020) were used for intravital imaging of cerebral microinfarction in mice brain *in vivo*. Briefly, cranial bone of mouse anesthetized by intraperitoneal injection of Zoletile (20 mg/kg) and Xylazine (10 mg/kg) mixture was exposed by skin incision and then the removal of connective tissue covering the cranial bone. Brain tissue of somatosensory cortex or posterior parietal association area was exposed by making a circular-shaped hole in the middle of parietal bone (typically 3 mm in diameter) with the dental drill (Strong 207A, Saeshin) under stereoscopic microscope. A 3 mm cover glass (64-0726, Warner Instruments) covered the exposed brain by using a fast-curing adhesive (Loctite 401. Henkel). To protect both of the incision area and the exposed cranial bone, the area except the cover glass was covered by using dental acrylic resin (1234, Lang Dental). After the imaging window implantation, mice were housed in cages for 3-4 weeks to allow a full recovery from the surgery before intravital imaging experiment. Mice showing normal brain vasculature with no sign of inflammation were selected and used for the intravital imaging and disease modeling.

### Intravital imaging of microinfarct

Mice were anesthetized by intraperitoneal injection of Zoletile (20 mg/kg) and Xylazine (10 mg/kg) mixture and mounted on a stereotaxic plate (US-R-10, Live Cell Instrument) with heating function to maintain the body temperature at 36.5 °C during intravital imaging. A cerebral microinfarction was induced for 3-5 seconds by 561nm laser illumination with 60X objective lens (LUMFLN60XW, NA 1.1, Olympus) after intravenous injection of 100μl Rose Bengal (15mg/ml; 330000, Sigma Aldrich)(Shih et al., 2013). Intravital imaging of ROS and cell death was performed at day 0 and 1 after injecting 250 μg of dihydroethidium (DHE, D11347, Invitrogen) and 75 μg of propidium iodide (PI, P4170, Sigma), respectively. Longitudinal imaging of Aldh1l1-GFP x NG2-DsRed mouse was performed for 30 days after injecting 30 μg of Alexa Flour 647-anti-CD31 antibody. Longitudinal imaging of NG2-DsRed x Kaede mouse was performed for 5 days after the injecting 400 μg of Alexa Flour 647-dextran (D22914, Thermo). Longitudinal imaging of vascular leakage in C57BL/6 mouse was performed for 5 days after injecting 600 μg of 3kDa TRITC-dextran (LIF-D-3308, Invitrogen) and 600 μg of 2MDa FITC-dextran (LIF-D-7137, Invitrogen). Time-lapse imaging of Aldh1l1-GFP mouse at day 1-1.5 was performed for 12 hr after injecting 30 μg of Alexa Flour 647-anti-CD31 antibody. Time-lapse imaging of NG2-DsRed x H2B-GFP mouse at day 3.5-4.5 was performed for 24 hr after injecting 30 μg of Alexa Flour 647-anti-CD31 antibody. Aldh1l1-GFP x NG2-DsRed x APP/PS1 mouse was imaged at 5.5 and 6.5 month-old mouse after injecting 600 μg of 3kDa TRITC-dextran (LIF-D-3308, Invitrogen), and 250 μg of Methoxy-X04 (4920, Tocris) for staining amyloid plaque.

### Immunofluorescence staining

Harvested brains were fixed with 4% paraformaldehyde (wt/vol, BPP-9016, Tech & Innovation) overnight at 4 °C and then sliced with Leica VT1200 vibrating microtome. Brain sections, 50μm thick, were permeabilized at 37 °C for 3 hr in 0.2% Triton X-100 (Sigma, T8787, Sigma) in phosphate-buffered saline (PBS, LB004, Welgene) and blocked overnight at 4 °C in 0.5% bovine serum albumin (BSA, A9418, Sigma) in permeabilization buffer. The sections were incubated for 3 days with primary antibodies in blocking buffer. Primary antibodies were diluted: NeuN (1:200, LIF-711054, Thermo), MAP2 (1:250, PA1-10005, Thermo), GLT-1 (1:100, AB1783, Merk), GFAP (1:200, AB5541, Merk) and LCN2 (1:200, AB2267, Merk). The sections were then washed for 1 day with permeabilization buffer and incubated for 1 day with secondary antibodies (all 1:1,000 dilution) in blocking buffer: goat anti-rabbit Alexa 555 (A21428, Thermo), goat anti-guinea pig Alexa 647 (A21450, Thermo), goat anti-chicken Alexa 555 (A21437, Thermo), goat anti-rabbit Alexa 647 (A21244, Thermo) and goat anti-chicken Alexa 647 (A32933, Thermo). After incubation, the sections were additionally stained by DAPI (D9542, Sigma) for 1 hr in PBS, and then mounted on slides. Images were obtained by custom-built confocal microscope system.

### Brain tissue clearing method

Cubic-L was prepared in deionized (DI) water with 10% (wt/vol) N-butyldiethanolamine (B0725, TCI) and 10% (wt/vol) Triton X-100 (T0818, Samchun). For tissue clearing, brain tissues were immersed and gently shaken in 5 mL Cubic-L solution in 37°C incubator for 1-2 days, exchanging the Cubic-L solution once every 24 hours. Once the white matters of the brain samples have been cleared, to minimize tissue damage, the tissue samples were immediately removed from the tissue clearing solution and washed in 5 mL of 0.1% (wt/vol) Triton X-100 in PBS (PBST) overnight. For samples that needed immunostaining, tissues were incubated in 37°C shaker with primary antibody, NeuN, in PBST, washed in PBST, incubated with secondary antibody, goat anti-rabbit Alexa 555, in PBST and washed with PBST at 37°C. For optical clearing, dPROTOS (RI=1.52) was prepared in DI water with 30% (vol/vol) 2,2’thiodiethanol (166782, Sigma), 30% (vol/vol) dimethyl sulfoxide (67-68-5, Samchun), and 106% (wt/vol) iohexol (RY2006001, Royal Pharm). Tissue samples were immersed in 5 mL dPROTOS and incubated at 37°C for 1 day to assure homogenous RI matching.

### Drug treatment

KDS2010, a MAO-B inhibitor, was dissolved at 2.5mg/ml in distilled water and orally administrated using oral zonde needle (JD-S-124, Jeungdo Bio & Plant) every day for 1 month (10 mg/kg daily) from day 1 after the induction of cerebral microinfarction.

### Statistical imaging analysis

To analyze alteration of tissue area, we constructed vasculature map on the bases of venule and arteriole branching points and then quantified tissue area following the onset of cerebral microinfarction. To quantify area of collagen deposition, maximal intensity projection (MIP) SHG images were obtained from 150 μm of z-stack images, and then measured area by using Image J software. Cell number, NeuN^+^ nuclear size and average Aldh1l1^+^ astrocyte size were measured from single field of view (FOV) images by spot analysis and surface rendering of a commercial image processing software, IMARIS (Bitplane). Pericyte processes lengths were quantified by using manual spot analysis of IMARIS. GFAP, Aldh1l1 and LCN2-positive area and co-localization area were measured by mean intensity using Image J software. To analyze the reactivity of individual astrocytes, we performed Sholl analysis, provided by Image J, and used region of interests (ROIs) from single Aldh1l1^+^ astrocyte images. Sholl analysis automatically measured by 4 μm of radius step size and then we quantified average process lengths and intersects in individual astrocytes. Mean intensity of FITC and TRITC was measured by using histogram of Image J software. Flow dynamics in NG2-DsRed x Kaede mouse were manually measured by tracing same capillaries for 5 days using Image J software, and quantified the ratio of dynamics according to flow, blockage, reflow and regression. Vascular area and lacunarity from 10 μm of MIP images were quantified by AngioTool software (version 0.6a). For statistical analysis, all quantification data were statistically analyzed by unpaired two-tailed t-test or one-way ANOVA with Tukey’s multiple comparison using Prism 5.0 (GraphPad), and statistical significance was set at p<0.05.

## Acknowledgements

We would like to thank Dr. W. Chung (Korea Advanced Institute of Science and Technology, Korea) for providing the Aldh1l1-GFP mouse, and Dr. Tomura and Dr. Miwa (Kyoto University, Japan) for providing the Kaede mouse. This work was supported by the Brain Research Program (2016M3C7A1913844) and the Basic Research Program (2020R1A2C3005694) funded by the Ministry of Science and ICT, Republic of Korea.

## Author Contributions

J.L. designed and performed the experiments, analyzed and interpreted data, and wrote manuscript. J.K., S.H., Y.K., S.A., R.K. performed experiments and interpreted data. H.C, K.P, Y.J., C.L., D.K., T.K. interpreted data, reviewed and edited the paper. P.K. supervised the study, design the experiments, interpreted data and wrote the manuscript. All authors read and approved the final manuscript.

## The Paper Explained

### PROBLEM

Cerebral microinfarcts, small ischemic lesions, contribute to the disruption of structural brain connection, which can progress to a cognitive impairment and dementia. However, little is known about the mechanistic link between microscopic cerebrovascular disruption and brain damage.

### RESULTS

We longitudinally observe a complex cellular-level changes of astrocyte, pericyte and vascular functions and integrity after microinfarct induction. Particularly, a fibrous collagen-rich glial scar formation is observed in the microinfarct with accumulation of GFAP+LCN2+ reactive astrocytes in the peri-infarct. Oral administration of KDS2010, a reversible inhibitor of monoamine oxidase B (MAO-B) in reactive astrocytes, significantly decreases the glial scar formation and astrocyte reactivity.

### IMPACT

Our longitudinal intravital imaging study of neurovascular pathophysiology in cerebral microinfarction demonstrate a susceptibility of astrocytes to ischemic brain injury and their reactive response leads to a long-term glial scar formation in cerebral cortex.

